# FGF2 promotes the expansion of parietal mesothelial progenitor pools and inhibits BMP4-mediated smooth muscle cell differentiation

**DOI:** 10.1101/2024.01.31.577512

**Authors:** Youngmin Hwang, Yuko Shimamura, Junichi Tanaka, Akihiro Miura, Anri Sawada, Hemanta Sarmah, Dai Shimizu, Yuri Kondo, Zurab Ninish, Kazuhiko Yamada, Munemasa Mori

## Abstract

Mesothelial cells, in the outermost layer of internal organs, are essential for both organ development and homeostasis. Although the parietal mesothelial cell is the primary origin of mesothelioma that may highjack developmental signaling, the signaling pathways that orchestrate developing parietal mesothelial progenitor cell (MPC) behaviors, such as MPC pool expansion, maturation, and differentiation, are poorly understood. To address it, we established a robust protocol for culturing WT1^+^ MPCs isolated from developing pig and mouse parietal thorax. Quantitative qPCR and immunostaining analyses revealed that BMP4 facilitated MPC differentiation into smooth muscle cells (SMCs). In contrast, FGF2 significantly promoted MPC progenitor pool expansion but blocked the SMC differentiation. BMP4 and FGF2 counterbalanced these effects, but FGF2 had the dominant impact in the long-term culture. A Wnt activator, CHIR99021, was pivotal in MPC maturation to CALB2^+^ mesothelial cells, while BMP4 or FGF2 was limited. Our results demonstrated central pathways critical for mesothelial cell behaviors.

## Introduction

The mesothelium, a distinctive cell type forming the pleural monolayer, envelopes the outermost layers of the viscera and facilitates the growth of developing organs. Despite the known fact that aberrant proliferation of adult mesothelial cells, often aggravated by asbestos exposure, can lead to mesothelioma through the manipulation of developmental pathways, the specific signaling processes that dictate progenitor pool expansion, embryonic mesothelial progenitor cell (MPC) maturation, and their differentiation into smooth muscle cells (SMC) remain poorly understood.

Anatomically, adult mature mesothelial cells of the parietal and visceral pleura encase the inner layer of the thorax and the outer layer of the lungs, respectively. Mouse lineage-tracing analyses showed that visceral mesothelial cells in developing lung pleura migrate inward and differentiate into vascular smooth muscle cells^1^, parabronchial smooth muscle cells^2^, and myofibroblast^3^, highlighting the multipotency of developmental MPCs. During development, the MPC arises from the exact origin, lateral plate mesoderm^4^, while mesothelioma tends to originate from parietal mesothelial cells^5^. Since carcinogenesis often hijacks developmental programs^6^, studying parietal mesothelial development could significantly advance mesothelioma diagnosis and treatment.

Mesothelioma, a rare and aggressive cancer often caused by carcinogens like asbestos or tar, has a notably high mortality rate^7^. The prevalence is high in the countries such as the United Kingdom, Australia, and New Zealand^8^. Various tumor markers were identified, including Calretinin (CALB2), mesothelin (MSLN), type III collagen (COL3A1), and secretory leukocyte peptidase inhibitor (SLP1)^9^. Despite the availability of treatments such as surgical decertification and chemotherapy, most cases are diagnosed at advanced stages, limiting effective intervention options ^10^. A better understanding of the behavior of MPCs in the parietal pleura during development could develop the prognostic markers of mesothelioma.

In mouse embryos, Wilms Tumor Protein 1 (WT1), a representative mesothelial cell marker, is expressed on visceral and parietal mesothelial cells from the lung and the thoracic cavity^1,11^. *WT1* knockout mice showed hypoplastic lung phenotype^11,12^ and the defects of human mesothelial cells by Congenital Diaphragmatic Hernia (CDH), also known to develop lung hypoplasia^13^.

Previous in vitro studies have shown that Fibroblast growth factor 2 (FGF2) and platelet-derived growth factor (PDGF) are required for the proliferation of adult mesothelial cells^14^. Notably, high expression of FGF2 in mesothelioma correlated with poor prognosis^15^.

Bone morphogenic protein 4 (BMP4) is expressed in the human adult peritoneal mesothelium and plays a pivotal role in mesothelial-to-mesenchyme transition (MMT), attenuating the TGF-beta-mediated MMT phenotype^16^. BMP4 is expressed ventral to the distal lung bud mesenchyme and at the distal lung bud tips of the endoderm^17,18^, but the association with the behavior of WT1^+^ MPC is unknown.

Additionally, Sonic hedgehog (SHH) and Retinoic acid (RA) are implicated in MPC migration and epithelial morphology transformation, respectively ^19^.

However, how these signaling pathways intertwine and distinctively regulate MPC pool expansion, differentiation, and maturation during development has yet to be determined, necessitating robust culture methods for detailed study.

This study successfully allowed us to establish the method to isolate and culture embryonic parietal MPC from developing pig and mouse thorax. By culturing these cells with a range of small molecules and growth factors, we aimed to elucidate the signaling pathways crucial for mesothelial cell development.

## Results

### Establishment of Cell Culture Protocol for the Expansion of Developing Pig Mesothelial Cells

The development of pig lungs undergoes embryonic, pseudo glandular, canalicular, and alveolar stages around embryonic day 19 (E19), E25, 60, and E90, respectively^20,21^. The developmental stage at which pig parietal mesothelial progenitor cells (MPCs) could be efficiently harvested was unknown. We harvested the parietal MPCs from the E80 canalicular stage thorax to have enough cell numbers.

To harvest a WT1^+^ developing MPC efficiently, we compared several methods previously reported^22–25^, including collecting pleural fluid, pinching porcine thoracic walls with tweezers, scaring it with scrapers, or trypsinizing the porcine thoracic wall. Among those methods, trypsinization with a 0.05% trypsin inside the E80 thoracic walls showed the highest yield of MPC collection (**Figure 1A**). Interestingly, 0.25% trypsin treatment to the thorax did not expand the MPC (**Figure S1A, B**). Previous papers showed the requirement of EGF for culturing EGF^23,25^. Contrary to expectations, MPC culture with EGF didn’t offer an apparent effect on MPC colony expansion (**Figure S1C**). To expand MPC efficiently, we coated the cell culture dish with extracellular matrix (ECM) molecules (type I collagen (Col I) and hyaluronic acid (HA)), given their expression in adult mature mesothelial cells^26,27^. We found that the isolated MPC showed the sustained expression of *Col I* expression and its receptor, *integrin beta 1* (*ITGB1*), but a relatively low expression of HA receptor (*CD44*) (**Figure 1B**). Indeed, Col I coating significantly enhanced MPC expansion compared to HA coating (HA) and an uncoated control (**Figure 1C, D**). Since the gelatin and Col1 share the integrin-binding motif, RGD sequence^28^, we cultured the MPC on the gelatin-coated dish and confirmed its efficacy in expanding MPCs^27^ (**Figure 1E**). Based on this, we performed all downstream analyses on the gelatin-coated dish. Additionally, we confirmed that mouse MPC can be collected and expanded well after the trypsinization directly on the E17.5 mouse canalicular ∼ sacculation stage thorax, noting that 0.25% trypsin was more effective for mouse MPCs than 0.05% (**Figure S1D-F**). These results underscore the robustness and effectiveness of our trypsinization-based protocol for isolating parietal MPCs in development.

**Figure 1.**
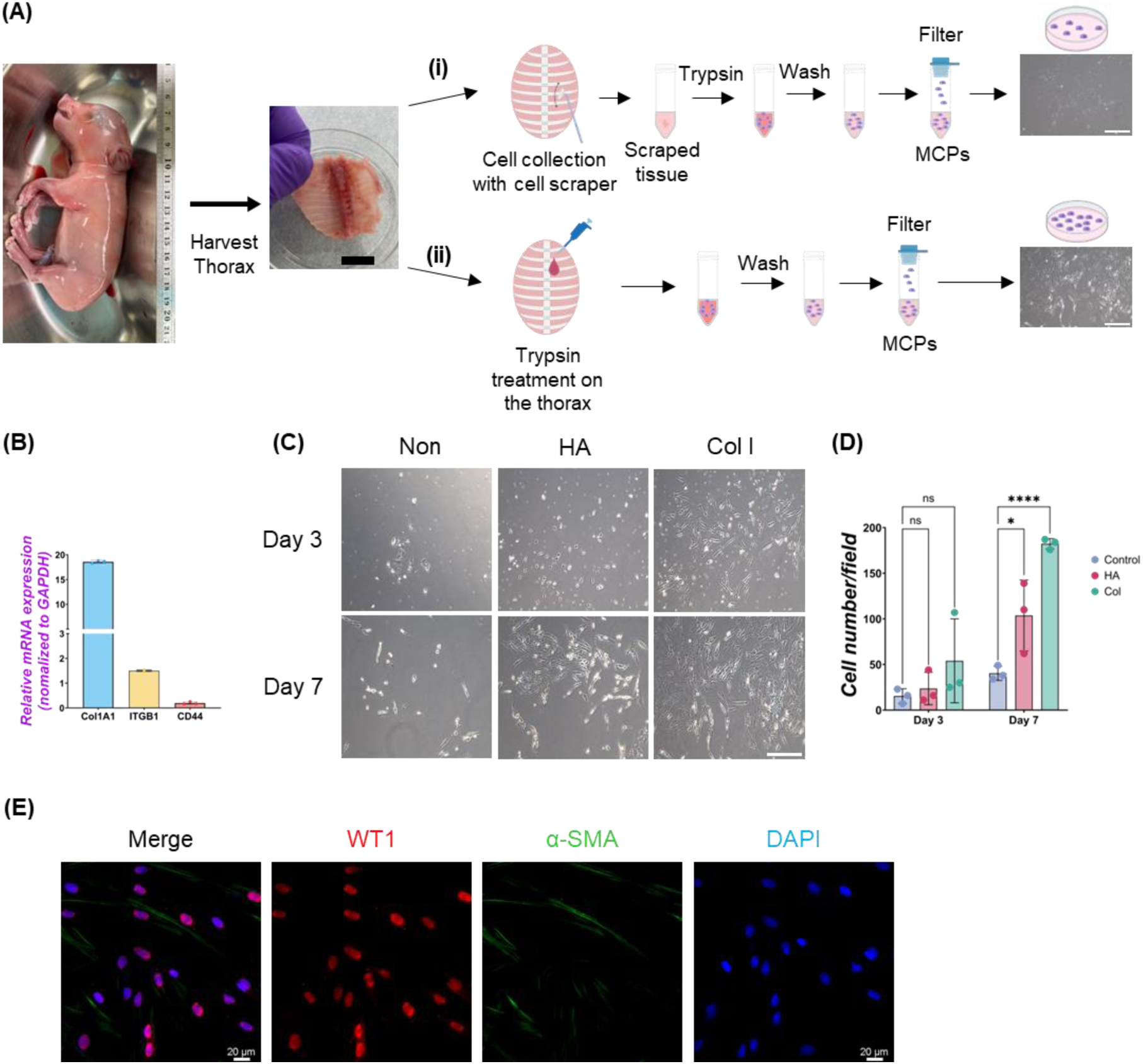
Isolation of Mesothelial cell progenitors (MPCs) from pig fetuses. (A) Schematic illustration of pig MPC isolation: The embryonic thorax (middle panel in A) was isolated from E80 pig fetuses (left panel in A) and treated with the following procedures. (i) scraping MPCs followed by trypsinization with 0.05% trypsin in the tube: (ii) trypsinization with 0.05% trypsin directly on the thorax. In both methods, the mesothelial cell was neutralized with DMEM + 10% FBS, followed by PBS washing and filtration with a cell strainer to remove the residual connective tissue. The trypsinization on the porcine thorax (ii) method showed a higher yield of MPC expansion than the scraping method (i) (right panels in A). (B) Graphs: quantitative qRT-PCR (RT-qPCR) analysis of type I collagen (*COL1A1*), integrin beta-1 (*ITGB1*), and *CD44* cultured in a basal culture medium. Error bars represent mean ± SD. Each plot showed different biological replicates (n = 3). Each gene expression was normalized with the housekeeping gene (*GAPDH*) expression. (C) Representative phase contrast images of MPCs isolated from E80 pig thorax cultured on different cell culture dish coating conditions. Col I: type I collagen coating, HA: hyaluronic acid coating, Non: non-coating. (D) Graphs: Quantification of the isolated pig MPC number per each field. Each plot showed different biological replicates (n = 3). (E) Representative immunofluorescence (IF) image of MPC after 3 days of culture. Red: WT1, Green: α-SMA, Blue: DAPI. Scale bars: (A) 1 cm, (C) 100 μm, (E) 20μm. *p<0.05, ****p<0.0001, ns: no significant difference by one-way ANOVA test and t-test in (D).

### FGF2 Promotes Expansion of Pig Mesothelial Progenitor Cells (MPCs)

While the role of FGF2 and PDGF in adult mesothelial cell proliferation is known, their impact during development is little known^14^. To confirm each molecule’s effect on developing MPCs, we cultured MPC with FGF2 and PDGF-BB for 3 days (**Figure 2**). PDGF-BB was chosen as the signaling molecule for the PDGF signaling pathway due to its binding potential to all PDGF receptors^29^. We found that FGF2 and PDGF-BB treatment increased total cell number as well as the WT1^+^ cell numbers compared to the basal condition control (**Figure 2A-D**). Ki67 immunostaining confirmed that FGF2 and PDGF-BB significantly increased proliferating cell numbers (**Figure 2A, B, E**). Notably, FGF2 and PDGF-BB induced a more than four times increase in proliferating Ki67^+^ WT1^+^ MPC proportion compared with the control in the short-term culture (**Figure 2F**). In contrast, the treatment with SU5402, a FGFR inhibitor, and CP 673451, a PDGFR inhibitor, significantly decreased both total and WT1 cell numbers (**Figure 2C, D**) by inducing 30∼40% of cell death, labeled by Cleaved Caspase3 (CASP3) 1-day post-treatment (**Figure S2**). These results suggested that the effect of endogenous FGF2 and PDGF activation cultured in the basal medium impacts ∼40% of MPC survival and that FGF2 and PDGF signaling may be essential for WT1^+^ MPC maintenance. To investigate the effect of FGF2 and PDGF on MPC pool expansion in the long term, we cultured the MPCs with FGF2 or PDGF-BB for 14 days and analyzed *WT1* mRNA expression by qPCR (**Figure 2G-I**). We found that FGF2 maintained *WT1* mRNA expression more than 5 times fold change compared to the control during long-term culture (**Figure 2H**), while the effect of PDGF-BB pool expansion did not significantly influence the *WT1* expression compared to the control over time (**Figure 2I**).

**Figure 2.**
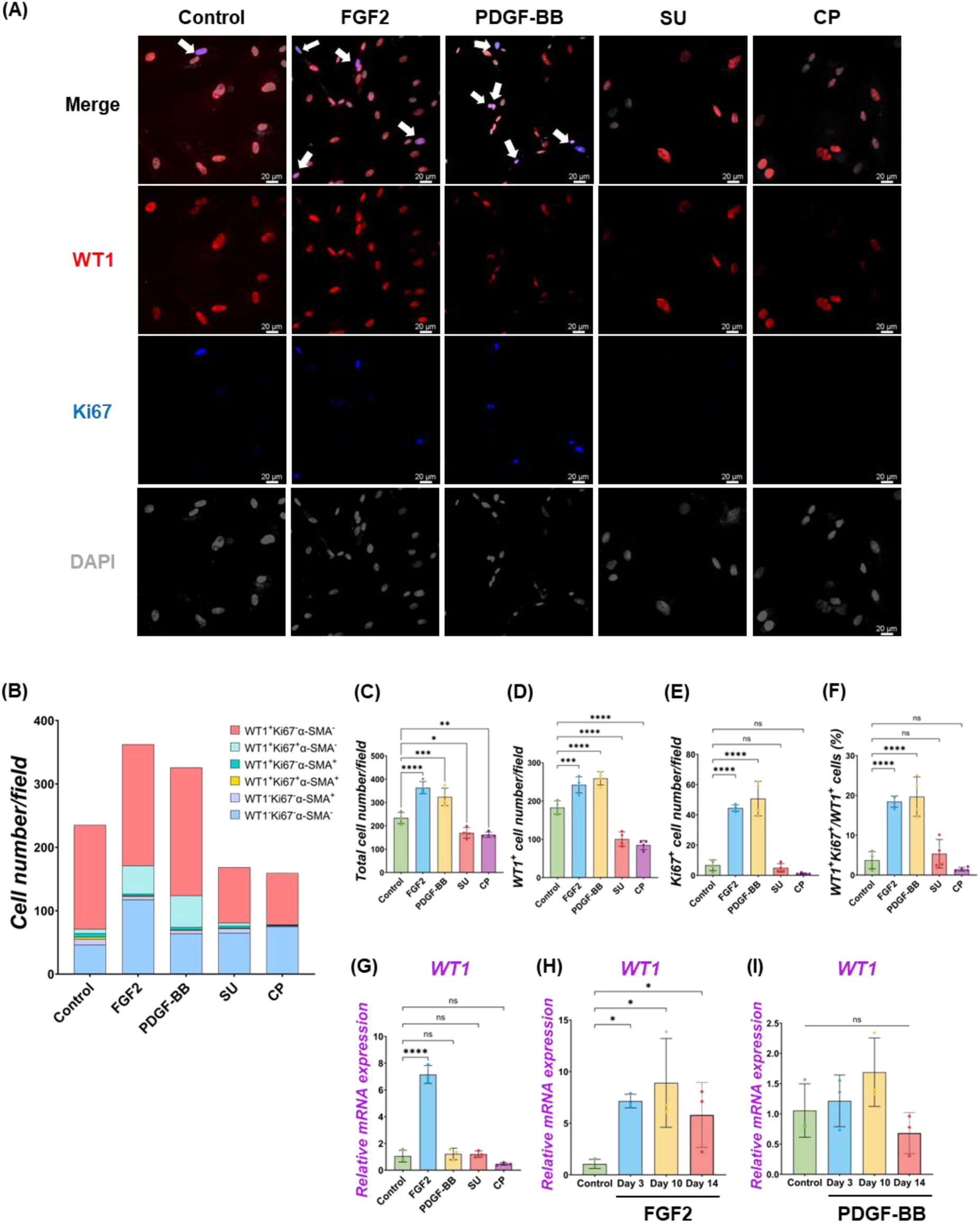
MPC self-renewal by FGF2, PDGF-BB stimulation. (A) Representative IF images of MPCs after 3 days of treatment with FGF2, PDGF, SU5404 (FGF signaling inhibitor, SU), a CP673451 (PDGF signaling inhibitor, CP), or Control (no treatment). FGF2 and PDGF-BB showed more cell numbers per field. WT1 (red), Ki67 (blue), DAPI (grey). Arrows (white): WT1^+^Ki67^+^ cells. (B) Graph: Quantification of cell numbers per field with each marker from IF images in (A). (n = 4) (C-F) Graphs: quantification of cell number from IF images with total cell number (C), WT1^+^ cell number (D), Ki67^+^ proliferative cell number (E), and proportion of WT1^+^Ki67^+^ proliferative MPCs (F). Error bars represent mean ± SD. Each plot showed different biological replicates (n = 4). (G-I) Graphs: RT-qPCR analysis of *WT1* mRNA expression after 3 days of culture with FGF2, PDGF-BB, SU, and CP (G). *WT1* mRNA expression during long-term culture by FGF2 (H) and PDGF-BB treatment (I). Error bars represent mean ± SD. Each plot showed different biological replicates (n = 3). Relative mRNA expression of each gene was normalized with the control basal culture condition. Scale bars = 20 μm. *p< 0.05, **p<0.01, ***p<0.001, ****p<0.0001, ns: no significant difference by one-way ANOVA test and t-test in (C-F).

These results suggest that FGF2 efficiently expands the MPC pools, but the PDGF-BB effect on the expansion is temporally and limited.

### BMP4 Drives Differentiation of MPCs into SMC

During the MPC control culture condition, WT1^-^α-SMA^+^ cells were observed (5.8 ± 3.3 %) (**Figure 2B**). We speculated that WT1^+^ MPCs could spontaneously differentiate into smooth muscle cells (SMCs), given that mouse visceral lung mesothelial cells differentiate into smooth muscle cells during mouse lung development^1,2^. To find which signaling molecules induce MPC differentiation into SMC, we cultured MPC with various small molecules and inhibitors with different concentrations and screened *α-SMA* mRNA expression by qPCR analysis (**Figure S3A**). We discovered that the BMP4 and ascorbic acid (AA) condition enhanced *α-SMA* mRNA expression compared to control among the tested conditions. Since BMP4 more dramatically induced SMC differentiation than AA, we focused on further analyses of BMP signaling. qPCR analyses found that BMP4 treatment showed significantly higher *α-SMA* mRNA induction both in short-term and long-term cultures, while BMP4 treatment had a transient effect on *WT1* mRNA increase only in the short term but did not sustain its impact in the long term (**Figure 3B, C**). In contrast, Dorsomorphin, a BMP4 inhibitor, significantly reduced *α-SMA* mRNA expression with no significant change in *WT1* mRNA expression (**Figure 3B**). Since the kinetics of *WT1* and *α-SMA* mRNA by BMP4 treatment indicated the MPC differentiation into SMC, we investigated the detailed cell fate change from MPC to SMC by immunostainings in short-term culture (**Figure 3A**). Consistent with the qPCR observations, immunostaining analysis showed a significantly increased α-SMA^+^ cell proportion (Control: 7.8 ± 1.7 % vs. BMP4: 31.4 ± 1.4 %) and the number by BMP4 treatment (**Figure 3A, D-F**), while dorsomorphin significantly reduced the α-SMA^+^ SMC proportion (6.7 ± 3.0 %). Unlike FGF2 and PDGF-BB (**Figure 2**), BMP4 treatment did not alter the total cell number, WT1^+^ MCP numbers, or WT1^+^ proportion but significantly increased Ki67^+^ cells (**Figure 3F-H**) while inducing about 20% of CASP3^+^ cell death, which might be the cell selection step (**Figure S2**). Indeed, BMP4 selectively eliminates the WT1^-^ Ki67^-^α-SMA^-^ unknown cell type while dorsomorphin significantly increased it (**Figure 3D**). Intriguingly, we observed a significantly increased proportion of WT1^+^α-SMA^+^ cells in WT1^+^ MPC (control: 9.9 ± 1.9 % vs. BMP4 group: 46.4 ± 6.4 %) by BMP4 treatment (**Figure 3I**), but proportion of WT1^-^α-SMA^+^ in SMC (control: 30.7 ± 15.4 % vs. BMP4 group: 43.7 ± 8.6 %) (**Figure 3J**) was not significantly changed. On the other hand, we did not observe any change in the proportion of WT1^+^α-SMA^+^ in α-SMA^+^ cells (**Figure 3K)**. These results indicate that BMP4 treatment primes the mesothelial progenitor pools to co-express WT1 and α-SMA, facilitating MPC differentiation into SMCs. Based on these results, including long-term culture, we concluded that the pivotal role of BMP4 is to induce parietal MPC differentiation into α-SMA^+^ SMC with losing WT1 expression.

**Figure 3.**
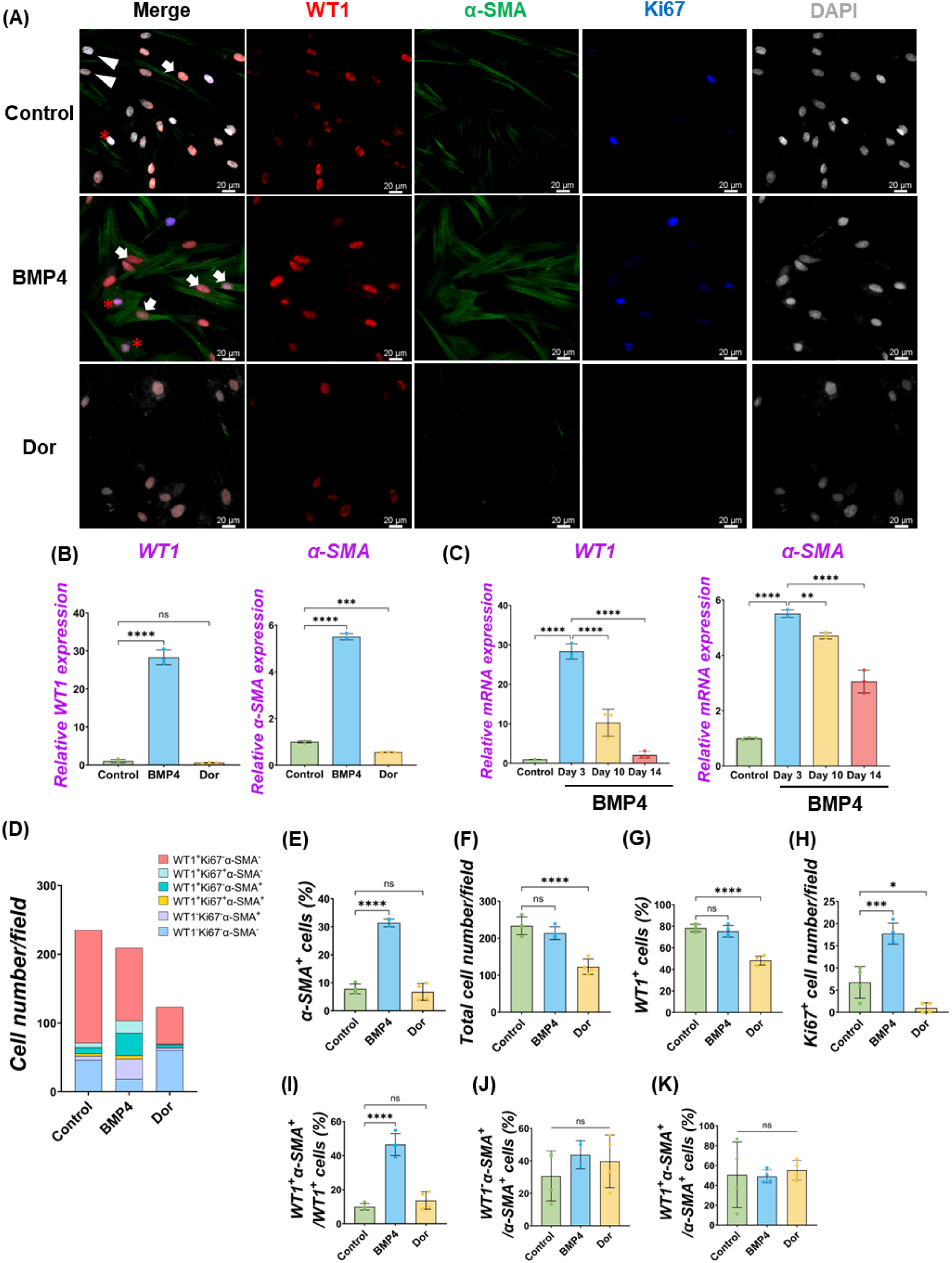
MPC differentiation into *α*-SMA^+^ smooth muscle cell by BMP4 stimulation. (A) Representative IF images of MPC after 3 days of treatment with BMP4, dorsomorphin (BMP signaling inhibitor, Dor), or Control (no treatment). BMP4 induced α-SMA expression, while a Dor reduced its expression. WT1 (red), α-SMA (green), Ki67 (blue), and DAPI (grey). Arrows (white): WT1^+^α-SMA^+^ cells, asterisks: WT1^+^Ki67^+^α-SMA^+^ cells, arrowhead (white): WT1^-^α-SMA^+^ cells. (B-C) Graphs: RT-qPCR analysis of *WT1* and *α-SMA* mRNA expression for 3 days of MPCs culture with BMP4, Dor, or Control (B) and long-term culture (C). Error bars represent mean ± SD. Each plot showed different biological replicates (n = 3). Relative mRNA expression of each gene was normalized with the control basal culture condition. (D) Quantification of cell numbers per field with each marker from IF images in (A). (E-I) Quantification of cell number from IF with α-SMA^+^ cell proportion (E), total cell number(F), WT1^+^ cell proportion (G), Ki67^+^ proliferating cell number (H), the proportion of WT1^+^α-SMA^+^ primed cells in WT1^+^ cells (I), WT1^-^α-SMA^+^ cells in SMA^+^ cells (J), and WT1^+^α-SMA^+^ cells in α-SMA^+^ cells (K). Error bars represent mean ± SD. Each plot showed different biological replicates (n = 4). Scale bars = 20 μm. *p< 0.05, **p<0.01, ***p<0.001, ****p<0.0001, ns: no significant difference by one-way ANOVA test and t-test in (B, C, E-K).

### FGF2 and PDGF-BB Suppressed MPC Differentiation into SMCs

We observed MPC progenitor pool regulation by FGF2 and PDGF-BB (**Figure 2**) and differentiation into α-SMA^+^ SMC by BMP4 (**Figure 3**), but it was unclear whether FGF2 and PDGF-BB influence the SMC pools. To address this, we performed qPCR analyses. We found that the decreased *α-SMA* mRNA expression by the FGF2 or PDGF-BB over time (**Figure 4A, B, S3**), and the further analysis of IF data showed that the proportion of α-SMA^+^ cells was significantly reduced by the FGF2 or PDGF-BB treatment (Control vs. FGF2 vs. PDGF-BB groups: 7.8 ± 1.7 % vs. 2.5 ± 0.5 % vs. 3.2 ± 0.4 %), while BMP4 significantly induced α-SMA^+^ cells (31.4 ± 1.4 %) (**Figure 4C**). In particular, PDGF-BB showed a dramatic decrease of *α-SMA* mRNA than FGF2 (**Figure 4B**). While there were no significant changes in the proportion of proliferating α-SMA^+^ cells, the proportion of WT1^+^α-SMA^+^ cells was significantly decreased by the FGF2 or PDGF treatment (Control vs. FGF2 vs. PDGF-BB groups: 9.9 ± 1.9 % vs. 3.8 ± 0.9 % vs. 3.4 ± 1.3 %) (**Figure 4D, E**). These results indicate that FGF2 and PDGF play a central role in MPC progenitor pool expansion by inhibiting the induction of WT1^+^α-SMA^+^ primed cells, leading to α-SMA^+^ smooth muscle cells (**Figure 4F**).

**Figure 4.**
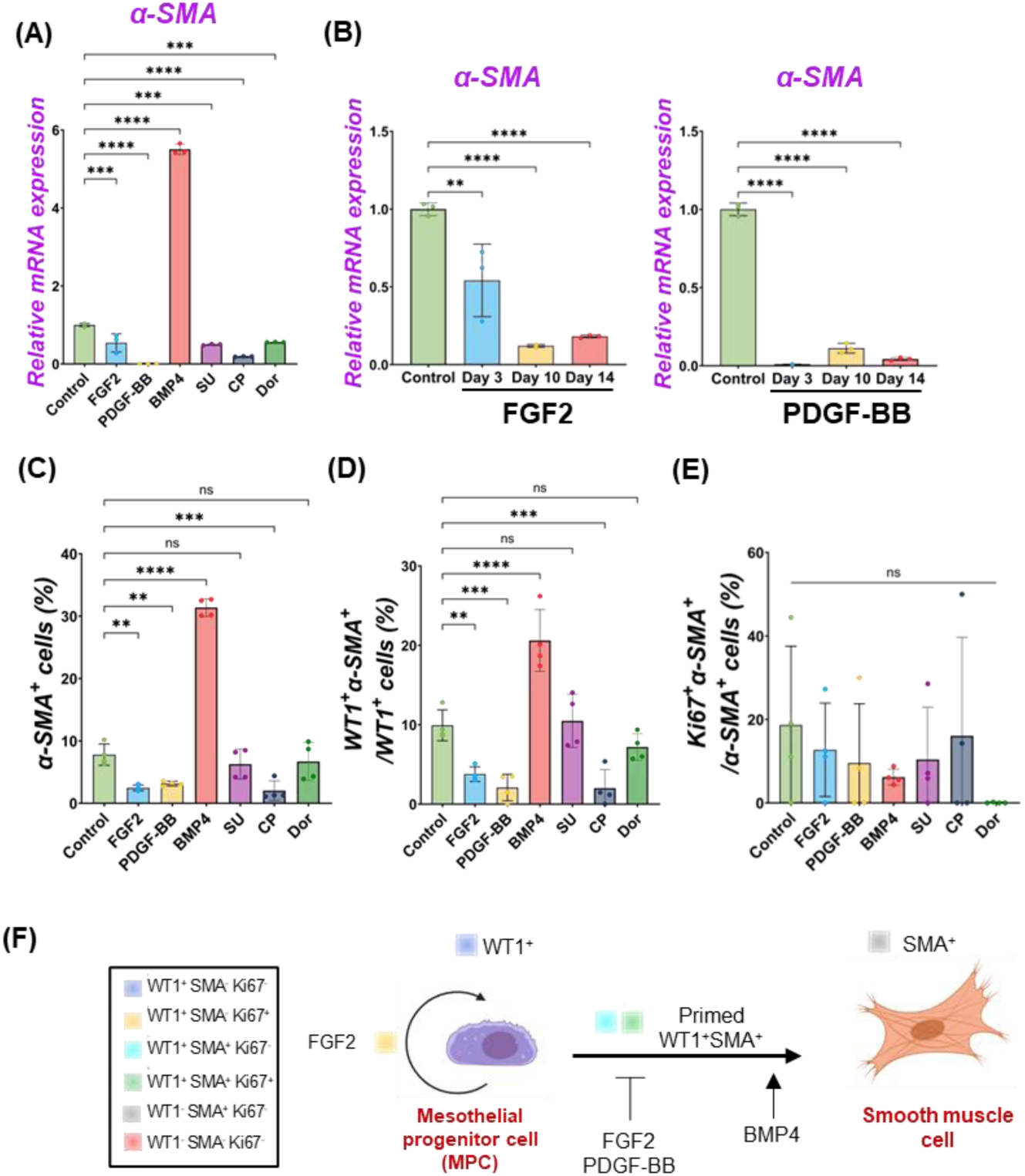
FGF2 and PDGF suppressed MPC differentiation into smooth muscle cells. (A-B) Graphs: RT-qPCR analysis of α-SMA. *α-SMA* mRNA expression after 3 days of MPCs culture with FGF2, PDGF-BB, BMP4, and its inhibitors (SU, CP, Dor) (A) and long-term culture of MPCs with FGF2, PDGF-BB (B). Error bars represent mean ± SD. Each plot showed different biological replicates (n = 3). Relative mRNA expression of each gene was normalized with the control basal culture condition. (C-E) Graphs: Quantification of cell proportion from IF of MPCs (from Figure 2**, 3**) with α-SMA^+^ cell proportion (C), proportion of WT1^+^α-SMA^+^ cells in WT1^+^ cells (D), and proportion of Ki67^+^α-SMA^+^ cells in α-SMA^+^ cells (E). Error bars represent mean ± SD. Each plot showed different biological replicates (n = 4) (F) Schematic summary of MPC self-renewal and differentiation into SMC by FGF2, PDGF-BB, and BMP4. **p<0.01, ***p<0.001, ****p<0.0001, ns: no significant difference by one-way ANOVA test and t-test in (A-E)

### Dominance of FGF2 Effect Over BMP Signaling in MPC Pool Regulation

Since we found FGF2 and PDGF suppressed BMP4-mediated MPC differentiation into SMC (**Figure 2-4**), we cultured MPC with the combination of FGF2 and BMP4 (FGF2 + BMP4) or PDGF-BB and BMP4 (PDGF-BB + BMP4) to investigate the potential counter effect. We found that MPC culture with FGF2 + BMP4 and PDGF-BB + BMP4 significantly suppressed the BMP4-mediated MPC differentiation into SMC with lower *α-SMA* mRNA expression than the BMP4 group (**Figure 5A)**. This mRNA expression trend was the same in the long-term culture (**Figure 5B**). Although the short-term treatment with FGF2 + BMP4 and PDGF-BB + BMP4 showed a decrease in *WT1* mRNA expression (**Figure 5A**), the long-term effect with FGF2 + BMP4 exhibited an increase in the *WT1* mRNA expression compared to controls (**Figure 5B**), consistent with the FGF2 effect (**Figure 2**). The long-term effect of PDGF-BB + BMP4 did not impact the *WT1* mRNA expression. Interestingly, the FGF2+BMP4 or PDGF-BB+BMP4 condition induced more cell proliferation with a higher total cell number than the BMP4 group in the short term (**Figure 5C-G**). In contrast, FGF2 + PDGF-BB and PDGF-BB + BMP4 conditions significantly increased WT1^+^ MPCs and proliferating cell numbers than the control condition in the short-term but could not sustain *WT1* mRNA expression in the long-term (**Figure 5A, E, F**). FGF2 + PDGF-BB and PDGF-BB + BMP4 conditions treatment significantly decreased α-SMA+ cells and showed no increase of primed WT1^+^α-SMA^+^ cells in WT1^+^ cells (**Figure 5G, H**). As we expected, there was no significant change in WT1^+^α-SMA^+^ cells in α-SMA^+^ cells (**Figure 5I**). These results suggest the critical role of FGF2 in maintaining the MPC pool and its self-renewal that counteracts the BMP signaling effects on MPC differentiation into SMC.

**Figure 5.**
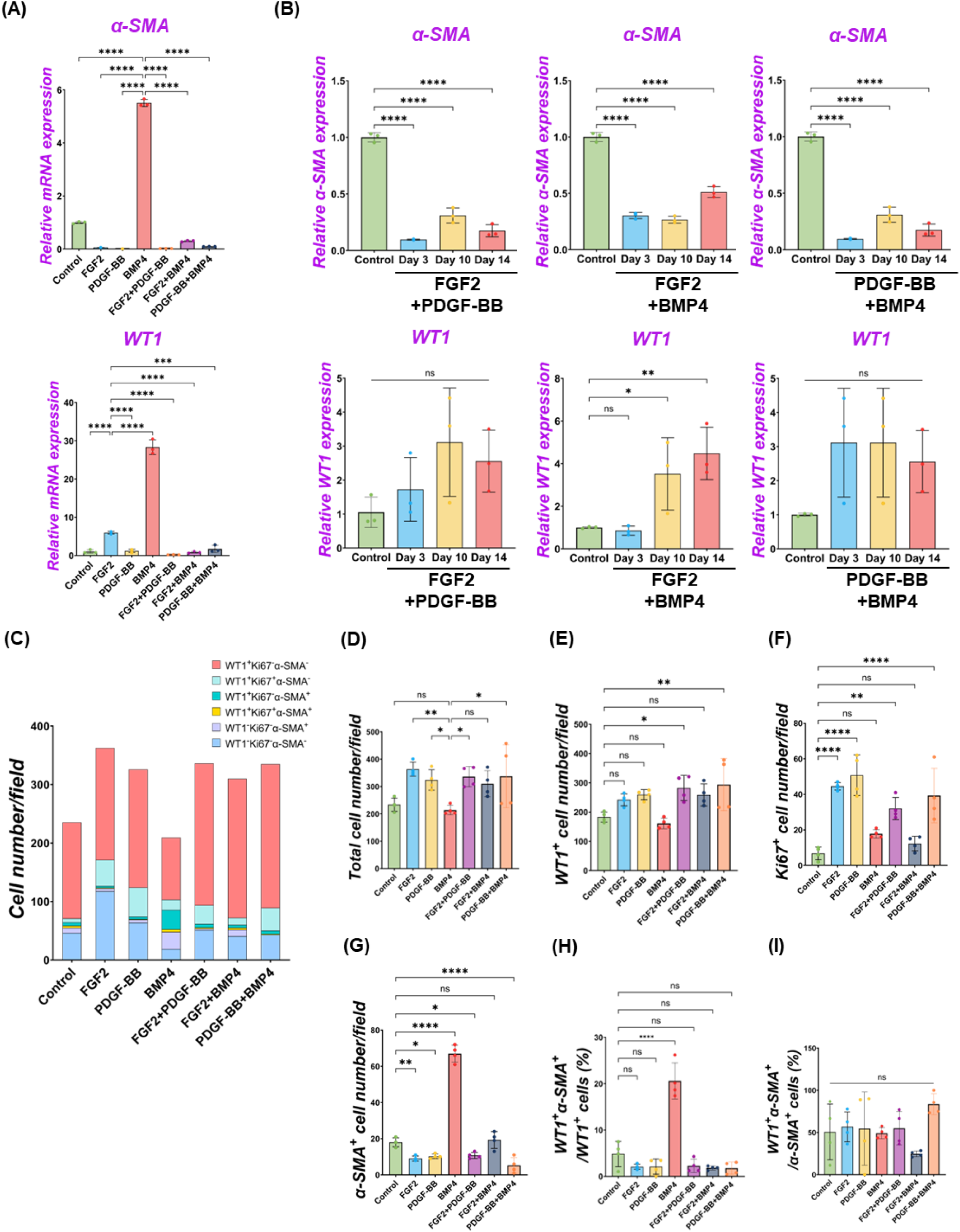
The dominance of FGF2 effect over BMP signaling in MPC pool regulation. (A-B) Graphs: RT-qPCR analysis of WT1 and *α-SMA* mRNA expression of MPC culture with signaling molecules and its combination during 3 days of culture (A) and long-term culture (B). (C) Graph: Quantification of cell numbers per field with each marker from IF images. (n = 4) (D-G) Graphs: quantification of cell number from IF with total cell number (D), WT1^+^ cells (E), Ki67^+^ cells (F), and α-SMA^+^ cells (G). (n = 4) (H-I) Graphs: proportion of WT1^+^α-SMA^+^ cells in WT1^+^ cells (H), proportion of WT1^+^α-SMA^+^ cells in α-SMA^+^ cells (I). Error bars represent mean ± SD. Each plot showed different biological replicates (n = 4) Scale bars = 20 μm. *p<0.05, **p<0.01, ****p<0.0001, ns: no significant difference by one-way ANOVA test and t-test in (A-I).

### Wnt Signaling Facilitates MPC Maturation

During development, mesenchymal β-catenin signaling controls parabronchial smooth muscle cell (PSMC) progenitors in the sub-mesothelial mesenchyme^2^. Wnt signaling is involved in the outer mesothelial pool size of the zebrafish swimbladder during development^28^. However, the molecular characterization of MPCs and their maturation during pig lung development have been little studied. To address this issue, we performed immunostaining of WT1 and CALB2 in pig and mouse lung development (**Figure S4**).

Developing porcine pleural mesothelial cells expressed high levels of WT1 in the E26 early pseudoglandular stage of porcine lungs, but the relative expression level in the peripheral layer of the lungs was decreased in the later stage (**Figure S4A, B**). In contrast, CALB2 expression was not detected in the peripheral layer of the lungs in the E26 and E40 early pseudoglandular stage but appeared in the canalicular stage and afterward (**Figure S4D, E**). These results indicate that CALB2 is the marker for mesothelial cell maturation during porcine lung development. We also confirmed that the WT1 expression pattern was also similar during mouse lung development, supported by previous studies^1,19^ **(Figure S4C)**, while CALB2 started to be expressed in the sub-peripheral layer from the E14.5 pseudoglandular stage in mouse lung development (**Figure S4F**).

To investigate the common MPC maturation markers across the species, we revisited the deposit single-cell RNA-seq (scRNA-seq) database of developing human^30^ and mouse^31^ lung mesenchyme (**Figure S5**). We found that *WT1* was highly expressed in the early pseudoglandular stage but decreased its expression in the late pseudoglandular and canalicular stages of human and mouse-developing lungs. *CALB2*, a mature mesothelial cell marker, was slightly observed but not abundant in human lung development. During mouse lung development, *CALB2* was observed in non-mesothelial cells. In contrast, mesothelin (*MSLN*) expression was observed in the late pseudoglandular stage of developing human lungs to the canalicular stage while around the E18 sacculation stage and afterward in the mouse lungs. These results suggest that decreased expression of *WT1* and increased *MSLN* are the evolutionarily conserved markers for MPC maturation, but *CALB2* is a pig-specific unique marker for MPC maturation. Based on these results, we examined pig MPC maturation in an in vitro study using *WT1*, *CALB2,* and *MSLN*.

We performed qPCR to screen the most potent signaling molecules regulating pig MPC maturation to CALB2^+^ and MSLN^+^ mature mesothelial cells (**Figure S3B, C**). Among them, we found that most signaling molecules induced the upregulation of *CALB2* and *MSLN* mRNA. In particular, the GSK3β inhibitor that acts as a Wnt activator (CHIR) showed the most dramatic increase in *CALB2* mRNA expression. Thus, we focused on analyzing Wnt signaling using CHIR in the MPC maturation. Three days of short-term CHIR treatment increased *WT1*, *CALB2*, and *MSLN* mRNA expressions, while the long-term CHIR treatment lost WT1^+^ MPC pools but relatively sustained *CALB2* expression (**Figure 6A**). Since high *WT1* mRNA expression is the landmark for immature MPC pool expansion, these results indicate that the MPC maturation by CHIR occurred as a long-term effect (**Figure 6A**).

**Figure 6.**
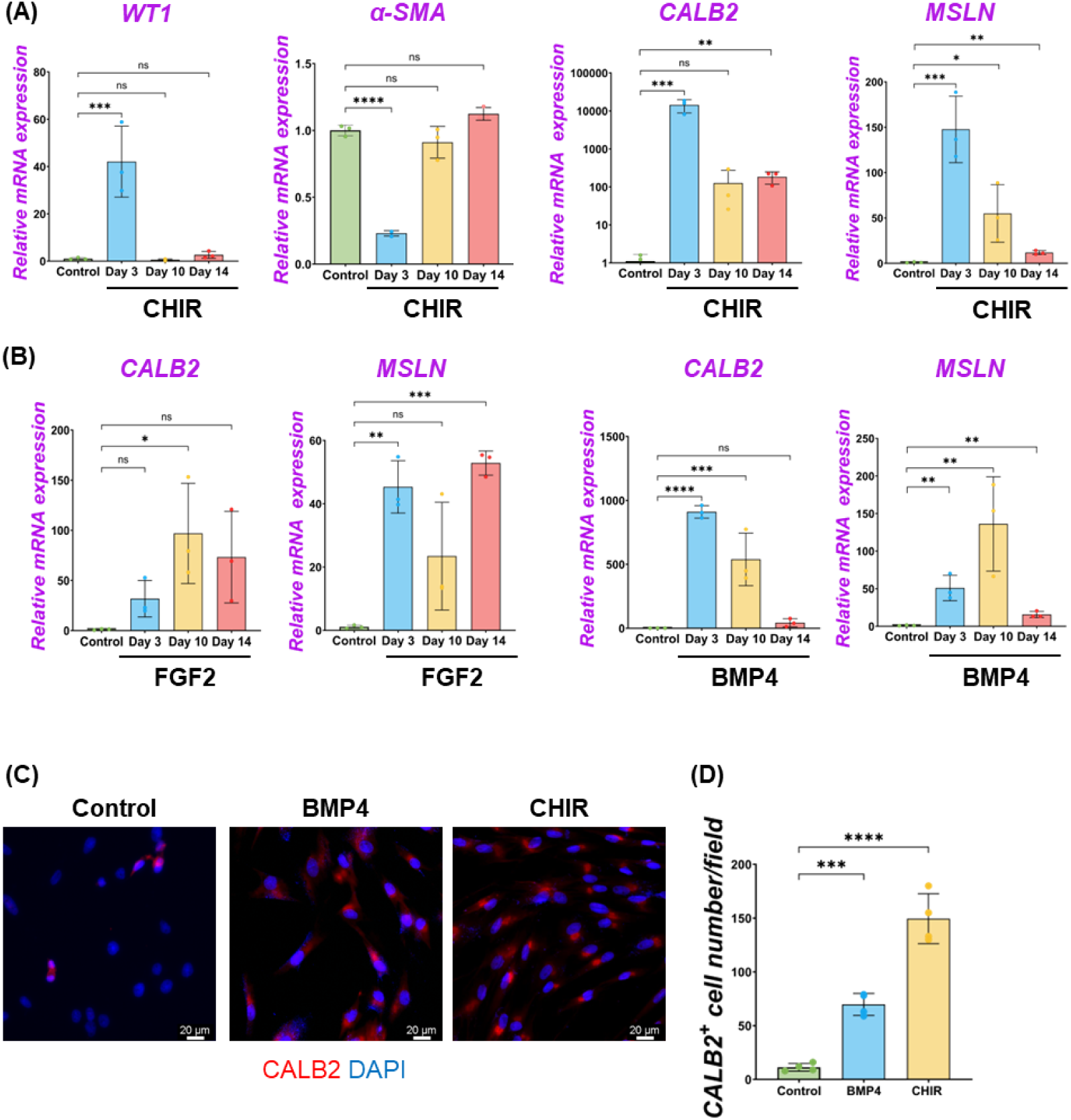
β-catenin (wnt) activation induced the maturation of MPCs to CALB2^+^ mature mesothelial cells. (A-B) Graphs: RT-qPCR analysis of *WT1*, *α-SMA*, *CALB2*, and *MSLN* mRNA expression for long-term culture of MPC treatment with CHIR99021 (CHIR) (A), and FGF2, BMP4 (B). Error bars represent mean ± SD. Each plot showed different biological replicates (n = 3). Relative mRNA expression of each gene was normalized with the control basal culture condition. (C) Representative IF images of MPCs after 3 days of treatment with BMP4 and CHIR. CALB2 (red), DAPI (blue). (C) Graph: quantification of CALB2^+^ cell number from IF. Error bars represent mean ± SD. Each plot showed different biological replicates (n = 4). Scale bars = 20 μm. *p<0.05, **p<0.01, ***p<0.001, ****p<0.0001, ns: no significant difference by one-way ANOVA test and t-test in (A, B, D).

Interestingly, we also found that long-term treatment with FGF2 or BMP4 significantly increased *MSLN* mRNA expression compared to the control (**Figure 6B**). However, FGF2 did not increase the mRNA expression of *MSLN* and *CALB2* in a dose-dependent manner in short-term culture, while BMP4 induced *CALB2* mRNA expression in a dose-dependent manner (**Figure S3C**). Furthermore, the *CALB2* mRNA upregulation by FGF2 or BMP4 was transient and relatively limited in the long-term treatment compared to the CHIR treatment (**Figure 6B**). Consistent with the qPCR results, the CALB2 immunostaining exhibited a consistent trend with qPCR results, indicating the increased CALB2^+^ cells by CHIR treatment (**Figure 6C, D**). As shown in the PDGF-BB effect, CHIR induced Ki67^+^ proliferative WT1^+^ cells and significantly increased total cell numbers compared to control (**Figure S6A-C**), while no WT1^+^ cell number or proportional change and reduced α-SMA^+^ cell number (**Figure S6D, E**). These results indicate that Wnt signaling activation induces MPC maturation into MSLN^+^ CALB2^+^ cells, corresponding to the expression pattern of CALB2 in porcine lung development.

## Discussion

Previous studies showed the markers of adult mesothelial cells or in mesothelioma, but it has been unclear how developing mesothelial progenitors shift the marker expressions and their association with cellular behaviors. We established an MPC expansion protocol that allows us to find the foundation of signaling pathways involved in MPC pool expansion, differentiation, and maturation. Technically, we could not expand the cells from the E40 or earlier time point’s thoracic wall in either method due to the low effectiveness of isolating MPCs even using swine specimens larger than mice (data not shown). Harvesting MPC exclusively from the lungs was also challenging because it contained various other cell types after the culture (data not shown). Based on these technical limitations, we focused on the MPC cellular analysis derived from the E80 thoracic walls. Of note, we also expand mouse MPC, in this culture condition, from the thorax at E17.0 ∼ E17.5 canalicular ∼ sacculation stage, corresponding to E80 pig developmental time points, indicating the robustness of our culture protocol to harvest and expand MPC (**Figure S1**).

FGF signaling pathways have been classically known as critical mitogens for both epithelium and mesenchyme^32–34^. Interestingly, mesothelial cells and mesothelioma have been characterized as epithelial-like and mesenchymal-like features^35,36^. We found that FGF2 has the most potent effect on MPC self-renewal in the long-term culture among tested conditions and inhibits BMP4-mediated SMC differentiation. Given that FGF2 high expression in mesothelioma is one of the critical prognosis factors and carcinogenesis often renders developmental program^37–39^, we speculate that targeting therapy for the FGF2 and its downstream, such as Spry2^40^, Ras^41^, or Sos^42^, may be critical for controlling FGF2^high+^ mesothelioma expansion and metastasis.

We found BMP4 signaling was critical for inducing MPC differentiation into SMC with an increase of α-SMA^+^ cells, including primed, transitioning WT1^+^α-SMA^+^ cells and differentiated WT1^-^α-SMA^+^ cells (**Figure 4**). The molecular mechanism of how BMP4 converts MPC to SMC needs to be determined in the future. Interestingly, our immunostaining analyses revealed that proliferating Ki67^+^α-SMA^+^ cells were never observed without tuning on WT1 (**Figure 4**). BMP4 initially induced WT1^+^Ki67^+^α-SMA^+^ transitioning cells but later lost the *WT1* mRNA expression (**Figure 3B**), suggesting that the critical role of BMP4 in MPC cell fate change to post-mitotic terminally differentiated SMC. Since retinoic acid treatment for acute leukemia patients induces terminally differentiated cells and is an effective therapy for those patients^43^, how BMP4 signaling activation would influence mesothelioma would be an attractive question.

Parietal MPC and lung peripheral MPC showed distinct morphology and function^44^. Our study showed that potential CALB2 descendants of MPC appeared around the neighboring WT1^+^ mesothelium (**Figure S4D, E**), supported by previous studies of mouse lung development^45^. There are remaining exciting questions regarding MPC maturation: about the role of CALB2 in porcine parietal MPC, its developmental distributions, how the parietal and lung-peripheral MPC distinctively mature, and how these MPC pools communicate during development. Interestingly, we did not observe CALB2^+^ cells on the parietal mesothelium during mouse development (**Figure S4F**). We examined three different antibodies against MSLN to investigate the maturation of MPC during development. However, MSLN expression was not detected in developing lungs and thorax, as in the previous study^19^, which is inconsistent with the scRNA-seq result (**Figure S5B**). This indicates that protein expression may be regulated at post-translational levels or require further technical advancements.

Interestingly, the WT1^+^ MPC showed α-SMA expression, reminiscent of porcine parietal mesothelial cells in the E26 early pseudoglandular stage (**Figure 1E, Figure S4A**), while it is uncommon in peripheral lung MPC. In our culture model, we used MPC at the canalicular ∼ sacculation stage. Our results indicate that porcine parietal MPCs may be a source of SMCs around the developing ribs.

We summarized MPC fate change by signaling molecules (**Figure 7**). Interestingly, FGF2 promoted the expansion of both WT1^+^ MPC and WT1^-^α-SMA^-^ pool compared to the control (**Figure 2B**). The WT1^-^α-SMA^-^ pool would involve CALB2^+^ mature mesothelial cells. However, BMP4 suppressed the WT1^-^α-SMA^-^ pool expansion (**Figure 3D**), while BMP4 also increased CALB2 expression in short-term culture (**Figure 6B, D**). This discrepancy suggests the existence of WT1^-^α-SMA^-^ CALB2^-^ unknown pool, which may have a role in the MPC regulation (**Figure 7**). Further analysis using genetic lineage tracing or single cell level bioinformatics analysis may reveal the lineage hierarchy, parietal MPC vs. peripheral lung MPC vs. WT1^-^α-SMA^-^ niche interactions, and association with mesothelioma, which will lead to further understanding of mesothelial development and pathogenesis.

**Figure 7.**
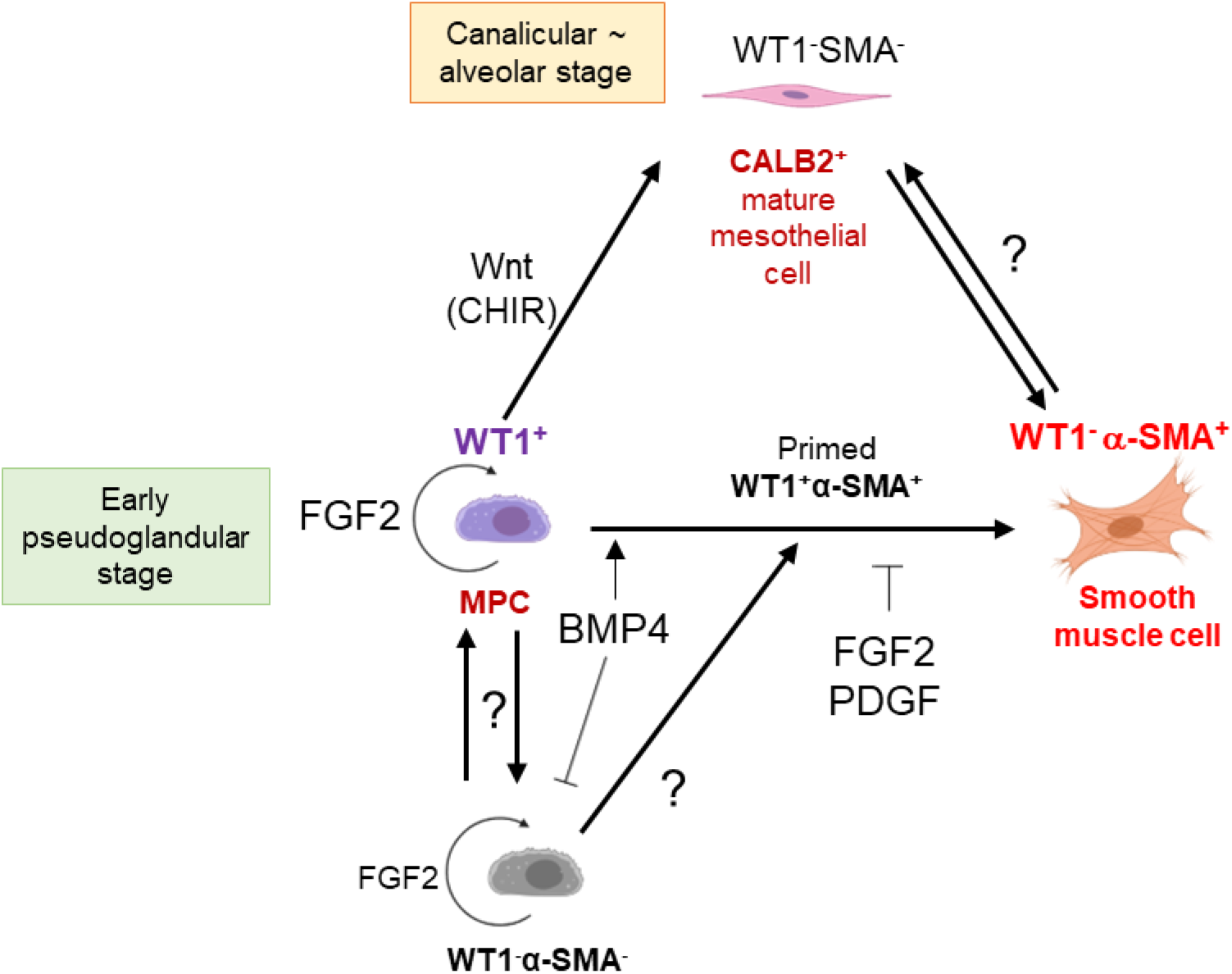
Schematic model of embryonic pig MPC cell behavior control by intertwined signaling. FGF2 induces self-renewal of WT1^+^ MPC. MPC differentiates into α-SMA^+^ SMC through primed WT1^+^α-SMA^+^ cells by BMP4 stimulation. FGF and PDGF signaling suppresses the BMP4-mediated SMC differentiation. Developing mesothelium shows stage-specific markers: high WT1 expression in the early pseudoglandular stage of porcine lung development and low WT1 expression and CALB2 expression in the calanlicular∼alveolar stage. Wnt activation by CHIR facilitates the MPC maturation process. The role of WT1^-^α-SMA^-^ unknown pools in MPC proliferation and differentiation is unclear.

## Acknowledgments

We thank Zurab Ninish for his technical assistance. We sincerely appreciate scientific input from Dr. Jianwen Que and Dr. Wellington Cardoso at the Columbia Center for Human Development (CCHD) and the members of Cardoso’s lab and CCHD. We acknowledge the support from the CCHD Medicine Microscopy Core (MMC) (NIH S10 OD032447-01). This work was funded by NIH-NHLBI 1R01 HL148223-01, DoD PR190557, PR191133 to M. M..

## Author contributions

Youngmin Hwang, Validation, Investigation, Visualization, Methodology, Writing – original draft; Yuko Shimamura, Junichi Tanaka, Akihiro Miura, Anri Sawada, Hemanta Sarmah, Dai Shimizu, Yuri Kondo, Investigation, Validation; Zurab Ninish, Kazuhiko Yamada, Methodology; Munemasa Mori, Conceptualization, Data curation, Supervision, Funding acquisition, Validation, Investigation, Methodology, Project administration, Writing – review and editing.

## STAR★Methods

### Key resources table

| REAGENT or RESOURCE | SOURCE | IDENTIFIER |
| --- | --- | --- |
| Antibodies |  |  |
| Rabbit anti-WT1 | Proteintech | Cat#12609-1-AP<br>RRID:AB_2216225 |
| Mouse anti-α-SMA | Bio-Rad | Cat#MCA5781GA<br>RRID:AB_3076452 |
| Chicken anti-Ki-67 | Novus Biologicals | Cat#NBP3-05538<br>RRID:AB_3076453 |
| Mouse anti-calretinin (2D7A9) | Thermo Fisher Scientific | Cat#66496<br>RRID:AB_2664066 |
| Chicken anti-calretinin | EnCor Biotechnology | Cat#CPCA-Calret<br>RRID:AB_2572241 |
| Rabbit anti-cleaved caspase-3 (Asp175) | Cell Signaling | Cat#9661<br>RRID:AB_2341188 |
| Rabbit anti-mesothelin (D9R5G) | Cell Signaling | Cat#99966<br>RRID:AB_2800323 |
| Rabbit anti-mesothelin (SP74) | Abcam | Cat#93620<br>RRID:AB_10563844 |
| Mouse anti-mesothelin (MSLN/2131) | Novus Biologicals | Cat#NBP2-79724<br>RRID:AB_3076454 |
| Donkey anti-mouse Alexa 488 | Invitrogen | Cat#A21202<br>RRID:AB_141607 |
| Donkey anti-mouse Alexa 647 | Invitrogen | Cat#A10042<br>RRID:AB_2534017 |
| Donkey anti-rabbit Alexa 568 | Invitrogen | Cat#A31571<br>RRID:AB_162542 |
| Donkey anti-chicken Alexa 488 | Jackson ImmunoResearch Labs | Cat#703-545-155<br>RRID:AB_2340375 |
| Goat anti-chicken HRP | Invitrogen | Cat#A16054<br>RRID:AB_2534727 |
| Chemicals, peptides, and recombinant proteins |  |  |
| Cy3 tyramide | AAT Bioquest | Cat#11065 |
| RBC Lysis Buffer (10x) | Biolegend | Cat#420301 |
| NucBlue™ Live ReadyProbes™ Reagent (Hoechst 33342) | Invitrogen | Cat#R37605 |
| rhEGF | R&D Systems | Cat#236-EG |
| rhFGF-basic | PeproTech | Cat#100-18B |
| SU5402 | MedChem Express | Cat#HY-10407 |
| rhPDGF-BB | R&D Systems | Cat#220-BB |
| CP673451 | MedChem Express | Cat#HY-12050 |
| rhBMP4 | R&D Systems | Cat#314-BP |
| Dorsomorphin | Tocris | Cat#3093 |
| CHIR99021 | MedChem Express | Cat#HY-10182 |
| Ascorbic acid | Fisher Chemical | Cat#FLA61100 |
| Retinoic acid | Sigma-Aldrich | Cat#R2625 |
| Purmorphamine | Tocris | Cat#4551 |
| Critical commercial assays |  |  |
| PrimeScript RT Master Mix (Perfect Real time) | Takara Bio | Cat#RR036B |
| Direct-zol™ RNA Purification kit | Zymo Research | Cat#R2062 |
| Luna Universal qPCR Master Mix | New England Biolabs (NEB) | Cat#M3003X |
| Deposited data |  |  |
| Human RNA-seq |  |  |
| Mouse RNA-seq |  |  |
| Pig RNA-seq |  |  |
| Experimental models: Organisms/strains |  |  |
| Mouse: Crl:CD1(ICR) | Charles River Laboratories | Strain: 022 |
| Yucatan pig | Sinclair BioResources | N/A |
| Oligonucleotides |  |  |
| qPCR primers, see Table S1 | This paper | N/A |
| Software and algorithms |  |  |
| GraphPad Prism 10.0 | https://www.graphpad.com/ | N/A |
| Cellpose | https://www.cellpose.org/ | N/A |
| ImageJ | https://imagej.net/ij/ | N/A |
| Leica Application Suite X (LAS X) | https://www.leica-microsystems.com/ | N/A |
| Other |  |  |
| Fetal Bovine Serum | Cytiva | Cat#SH30088.03HI |
| Trypsin-EDTA (0.05%) | Gibco | Cat#25300054 |
| Trypsin-EDTA (0.25%) | Gibco | Cat#15050065 |
| DMEM medium, high glucose | Cytiva | Cat#SH30243.02 |

### Resource availability

#### Lead contact

Further information and requests for resources and reagents should be directed to and will be fulfilled by the lead contact, Munemasa Mori.

#### Materials availability

All biological materials used in this study are available from the lead contact upon request.

#### Data and code availability

- This paper does not report original code.
- Any additional information required to reanalyze the data reported in this paper is available from the <u>lead contact</u> upon request.

### Experimental model and study participant details

#### Animals

All surgical procedures were conducted under the approval of the Columbia University Institutional Animal Care and Use Committee and USAMRMC Animal Care and Use Review Office (ACURO). For pig experiment, Timed-pregnant Yucatan miniature sows were obtained from Sinclair BioResources. For mouse experiment, CD-1 mice (male (8 weeks), female (8 weeks)) were purchased from Charles River Laboratories.

#### Parietal pig mesothelial progenitor cell (MPC) isolation

E80 Yucatan pig embryo was surgically collected from the Yucatan pig mother. After euthanization, the thorax was collected. For MPC isolation, we performed 2 methods; 1) the mesothelial tissue was isolated from the E80 pig thoracic wall with a cell scraper (Fisher Scientific), by following incubation in 0.25% trypsin-EDTA solution for 20 min at 37 °C and 2) 0.25% trypsin treatment on the thoracic wall, by following 20 min incubation at 37 °C. After trypsin-EDTA treatment, the dissociated cell was washed with PBS by centrifuge and replacement of the PBS (350 x g, 5 min, 4 °C). The cell pellet was incubated in RBC lysis buffer solution for 10 min at 4°C for RBC lysis (Biolegend), following PBS wash by centrifuge (350 x g, 5 min, 4 °C). After washing with PBS, the cell pellet was filtered with a cell strainer (40um pore size, MTC Bio) and seeded on a type I collagen (from rat tail, Sigma-Aldrich)-coated 6-well tissue culture plate. The MPCs (P0) were cultured in MPC culture medium (DMEM (high glucose, Gibco) + 10% FBS (Cytiva) + 1% pen/strep (Gibco)) for 7 days. For passage, MPCs were washed with PBS and dissociated with 0.05% trypsin-EDTA (Gibco) for 5min at 37°C). For MPC culture and its analysis for the experiments, passages 6-8 MPC were cultured on gelatin-coated tissue culture plates.

#### Parietal mouse mesothelial progenitor cell (MPC) isolation

Mouse MPC was isolated from E17.5 embryonic thorax by treatment of 0.05 % or 0.25 % trypsin-EDTA (Gibco) solution for 20 min at 37 °C. The isolation procedure was the same as pig MPC isolation. The mouse MPC was cultured in an MPC culture medium with the replacement of the cell culture media every other day.

#### Parietal pig mesothelial progenitor cell (MPC) culture

To investigate the MPC cell fate by signaling molecules, MPCs were cultured in the MPC culture medium with various signaling molecules (FGF2 (Peprotech), PDGF-BB, BMP4 (R&D systems), retinoic acid (RA, Sigma-Aldrich), CHIR99021 (MedChem Express), ascorbic acid (AA, Fisher Chemical), purmorphamine (Shh, Tocris)) and the inhibitors (SU5402 as FGFR inhibitor (MedChem Express), CP673451 as PDGFR inhibitor (MedChem Express), and dorsomorphin (Tocris) for 3, 10, or 14 days. During MPC culture, the MPC culture medium, including signaling molecules, was replaced every other day and passaged at day 3, 6, and 10 to avoid full confluency.

#### RT-qPCR

mRNA was isolated from MPCs with Direct-zol RNA Microprep isolation kit (Zymo Research) after lysis of MPCs with IBI isolate total reagent (IBI Scientific). For cDNA synthesis, the isolated mRNA was mixed with PrimeScript RT Master Mix (Takara bio), followed by cDNA synthesis protocol. For RT-qPCR analysis, the synthesized cDNA was mixed with qPCR primers and Luna Universal qPCR Master Mix (New England Biolabs (NEB). RT-qPCR was conducted with Quantstudio (Applied Biosystems). mRNA expression of each gene was normalized with the housekeeping gene (GAPDH). The relative mRNA expression of the genes was normalized with the control group (MPC culture in DMEM + 10% FBS + 1% pen/strep).

#### Immunofluorescence (IF)

For cell sample preparation, MPCs were fixed with 3.7% paraformaldehyde (PFA) for 10 min at room temperature. For tissue sample preparation, 10um-frozen sectioned tissue samples were washed with PBS 3 times, followed by antigen retrieval with citrate-based buffer (Vector Laboratories) in the microwave for 8 min. After washing the cells and the tissue samples with PBS 3 times, the primary antibodies in dilution solution (0.25% triton X-100 + 0.75% BSA in PBS) were treated to the samples and incubated at 4°C for overnight.

After 3 times PBS wash on the following day, the secondary antibodies and DAPI were treated (0.75% BSA in PBS) for 1 hour at room temperature. Then, the sample was mounted with a coverglass, anti-fade reagent (Invitrogen). For pig cell/tissue CALB2 staining, primary antibody-treated samples were treated with HRP conjugated anti-chicken antibody (in PBS) and incubated for 30 min at room temperature. After PBS wash, Cy3 tyramide (1:1000 diluted in 100 mM borate + 0.1% Tween-20 + 0.003 % H_2_O_2_ solution (pH 8.5)) was treated in the samples and incubated for 15 min at room temperature in the dark. After PBS wash, the samples were mounted with a coverglass and an anti-fade reagent (Invitrogen).

The cell samples were visualized with a Leica DMI microscope (Leica). The tissue samples were visualized with a Zeiss confocal microscope (Zeiss).

#### RNA-seq data analysis

For human and mouse RNA-seq data analysis, we utilized the database from the previou*s studies.*^30,31^

### Quantification and statistical analysis

Quantification of cell number in the phase contrast images was conducted by ImageJ. For immunostained cell (single-immunostained and co-immunostained cell population) and DAPI-stained cell counting from IF images, Cellpose software was used. The mean fluorescence intensity (MFI) of each IF sample was measured in the non-overlapping random fields using ImageJ software. Data analysis was performed using Prism 10. Data acquired by performing biological replicas ((n = 3) for RT-qPCR and phase contrast images, (n = 4) for

IF images) of three or four independent experiments are presented as the mean ± standard derivation (SD). Statistical significance was determined using a one-way ANOVA or a two-tailed t-test. *p < 0.05, **p < 0.01, ***p < 0.001, ****p < 0.0001, ns: non-significant.

### Additional resources

Human scRNA-seq: https://cellxgene.cziscience.com/e/f9846bb4-784d-4582-92c1-3f279e4c6f0c.cxg/

Mouse sdRNA-seq: https://lungcells.app.vumc.org/

