## Supplemental Information for "FGF2 promotes the expansion of parietal mesothelial progenitor pools and inhibits BMP4-mediated smooth muscle cell differentiation"

**Figure S1.** **Comparison of MPC isolation and culture conditions.** (A) Representative phase contrast images of pig MPCs isolated from E80 pig thorax with 0.25% trypsin treatment. 0.25% trypsin-treated MPC did not expand well in either of the conditions. (B) Graph: quantification of the isolated MPC number per field from phase contrast images in (A). Col I: type I collagen coating, HA: hyaluronic acid coating, Non: non-coating. Error bars represent mean ± SD. Each plot showed different biological replicates (n = 3). (C) Phase contrast images of the MPCs cultured on a type I collagen-coated well plate with hEGF (10ng/ml) or without hEGF. (D-F) MPC isolation from E17.5 mouse embryo. (D) Scheme of mouse MPC isolation by trypsin treatment on E17.5 mouse thoracic walls. (E) Representative phase contrast images of the mouse MPCs cultured on type I collagen-coated cell culture plates. (F) Graph: quantification of cell number per field from (E). Error bars represent mean ± SD. Each plot showed different biological replicates (n = 3). Scale bars = 100 μm. *p<0.05, ns: no significant difference by one-way ANOVA test and t-test in (B, F).

**Figure S2.** **Cell death induction** **in MPCs with each signaling molecule.** (A) Representative IF images of MPC 24 hr of culture with the treatment of FGF2, PDGF-BB, BMP4, SU, CP, or Dor. Red: Cleaved Caspase3 (CASP3), green: α-SMA, blue: DAPI. Scale bars = 20 μm. (B-C) Quantification of CASP3^+^ cell proportion (C) and α-SMA^+^CASP3^+^ cells in CASP3^+^ cells per field (C) from IF images in (A). Error bars represent mean ± SD. Each plot showed different biological replicates (n = 4). *p<0.05, ***p<0.001, ****p<0.0001, ns: no significant difference by one-way ANOVA test and t-test in (B, C).

**Figure S3.** **Gene expression analysis by RT-qPCR analysis of MPC cultures using various signaling molecules.** (A-D) Graph: RT-qPCR analysis of *α-SMA* (A), *CALB2* (B), and MSLN (C) mRNA expressions after 3 days of MPCs with various signaling molecules. Error bars represent mean ± SD. Each plot showed different biological replicates (n = 3). Relative mRNA expression of each gene was normalized with the control basal culture control condition. RA: retinoic Acid, PMA: Purmorphamine, AA: Ascorbic Acid.

**Figure S4. Expression of embryonic mesothelial cell markers in developing pig and mouse lungs.** (A, C, D, F) Representative IF images of the peripheral lung regions in pig (A) or mouse (C) lung development using WT1 (A, C) or CALB2 (D, F) antibody. Green: WT1, red: α-SMA (A, C), CALB2 (D, F), blue: DAPI. Th: Throax, Lu: Lung (B) Graph: mean fluorescence intensity (MFI) of WT1^+^ area on the peripheral region (mesothelial layer) in the peripheral layer of lungs (the outer layer of dotted line) per field. Error bars represent mean ± SD. Each plot showed different biological replicates (n = 4). (E) Graph: MFI of CALB2^+^ area (peripheral: the MFI in the outer layer of dotted line; or non-peripheral region: the MFI in the inner layer of dotted line, top panel), peripheral (bottom left panel), and non-peripheral (bottom right panel) region per field from IF images in (D). Error bars represent mean ± SD. Each plot showed different biological replicates (n = 4). (F) Representative IF images of mouse embryos. Red: CALB2, blue: DAPI. Scale bars = 20 μm. *p<0.05, **p<0.01, ***p<0.001, ****p<0.0001, ns: no significant difference by one-way ANOVA test and t-test in (B, E).

**Figure S5.** **Single-cell RNA-seq (scRNAseq) analysis of mouse or human lung mesothelial cell development.** (A) UMAP feature plots of scRNAseq of human embryonic lung mesenchyme (from a scRNAseq database (He et al., 2022, Cell))^30^, and (B) mouse embryonic lung (from a scRNAseq database (Negretti et al., Development))^31^ highlighting the expression of *WT1, MSLN, CALB2* in the developing mesothelial cells during human or mouse lung development. Staging is corresponding to the original article analyses^30,31^. (A) pcw: post-conception weeks, red rectangle in (A, B): mesothelial cells regions. Right panels in (A, B): Enlarged images of the red rectangle area. Pseudo: pseudoglandular stage, Canalicular: canalicular stage, alveolar: alveolar stage.

**Figure S6. Effect of Wnt signaling on MPC differentiation into SMC.** (A) Representative IF images of MPC after 3 days of CHIR treatment. Red: WT1, blue: Ki67, green: α-SMA, grey: DAPI. (B) Graph: quantification of cell numbers per field with each marker from IF images in (A). (C-F) Graphs: quantification of cell number from IF images with total cell number (C), WT1^+^ cell number (D), Ki67^+^ cell number (E), α-SMA^+^ cell number (F). Error bars represent mean ± SD. Each plot showed different biological replicates (n = 4). Scale bars = 20 μm. **p<0.01, ns: no significant difference by t-test in (B-F).

**Table S1. The list of RT-qPCR primer sequences used in this study.**

**Figure S1.**

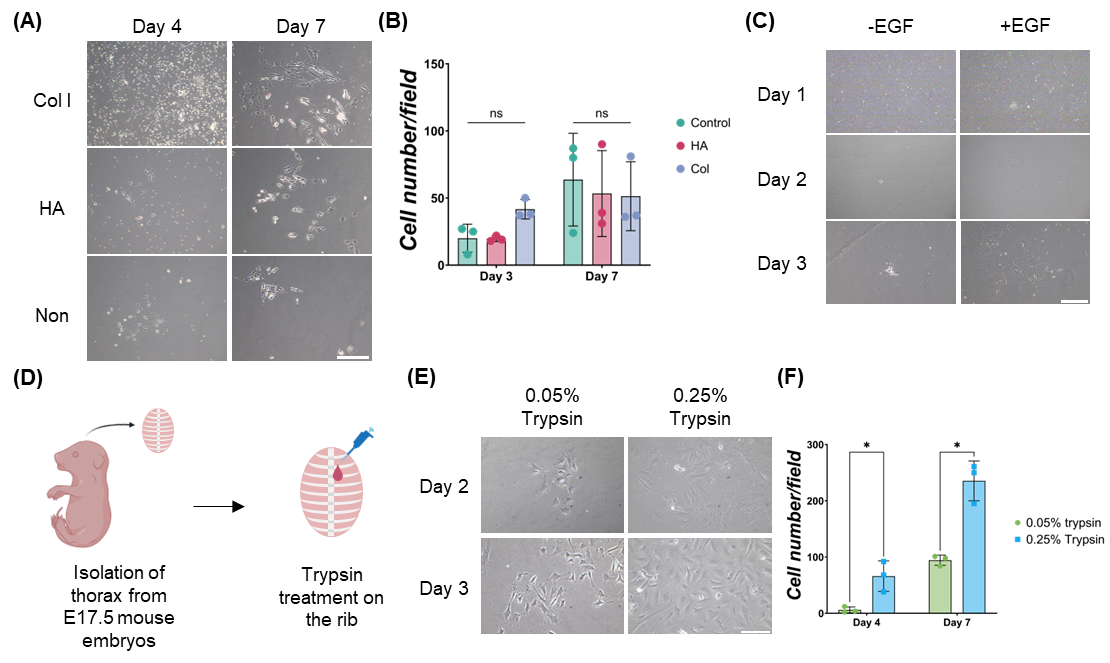

**Figure S2.**

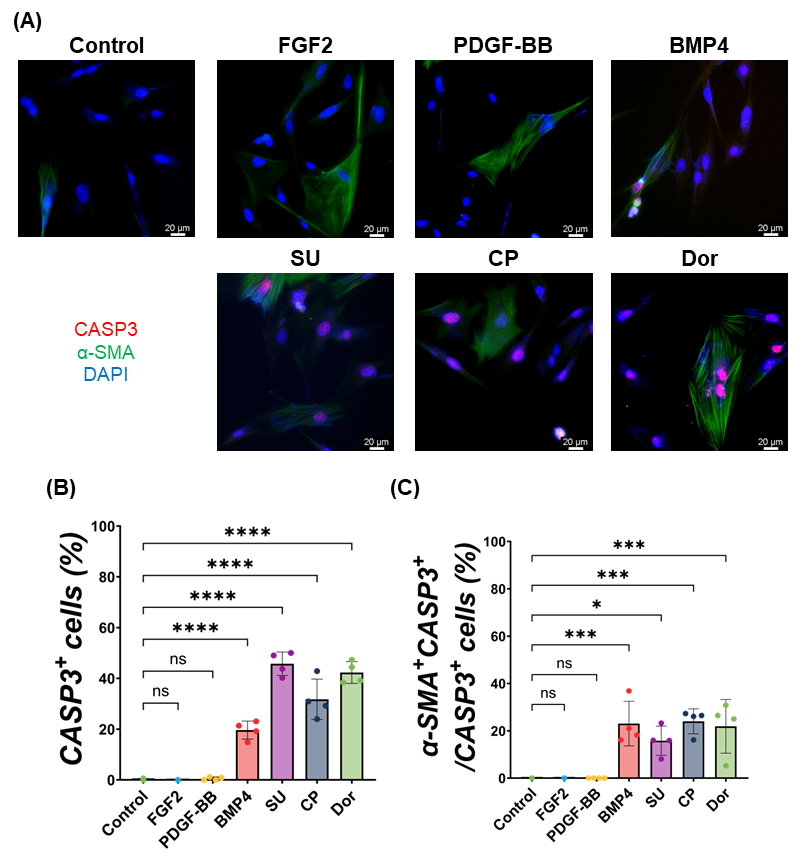

**Figure S3.**

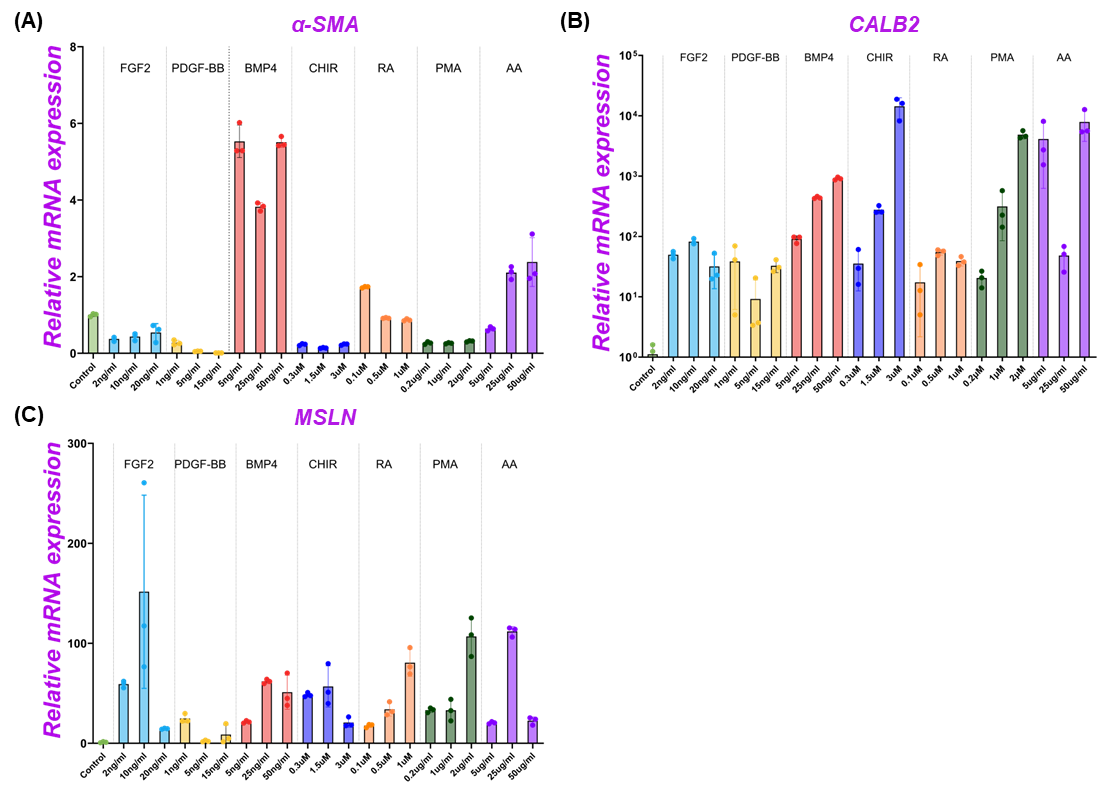

**Figure S4.**

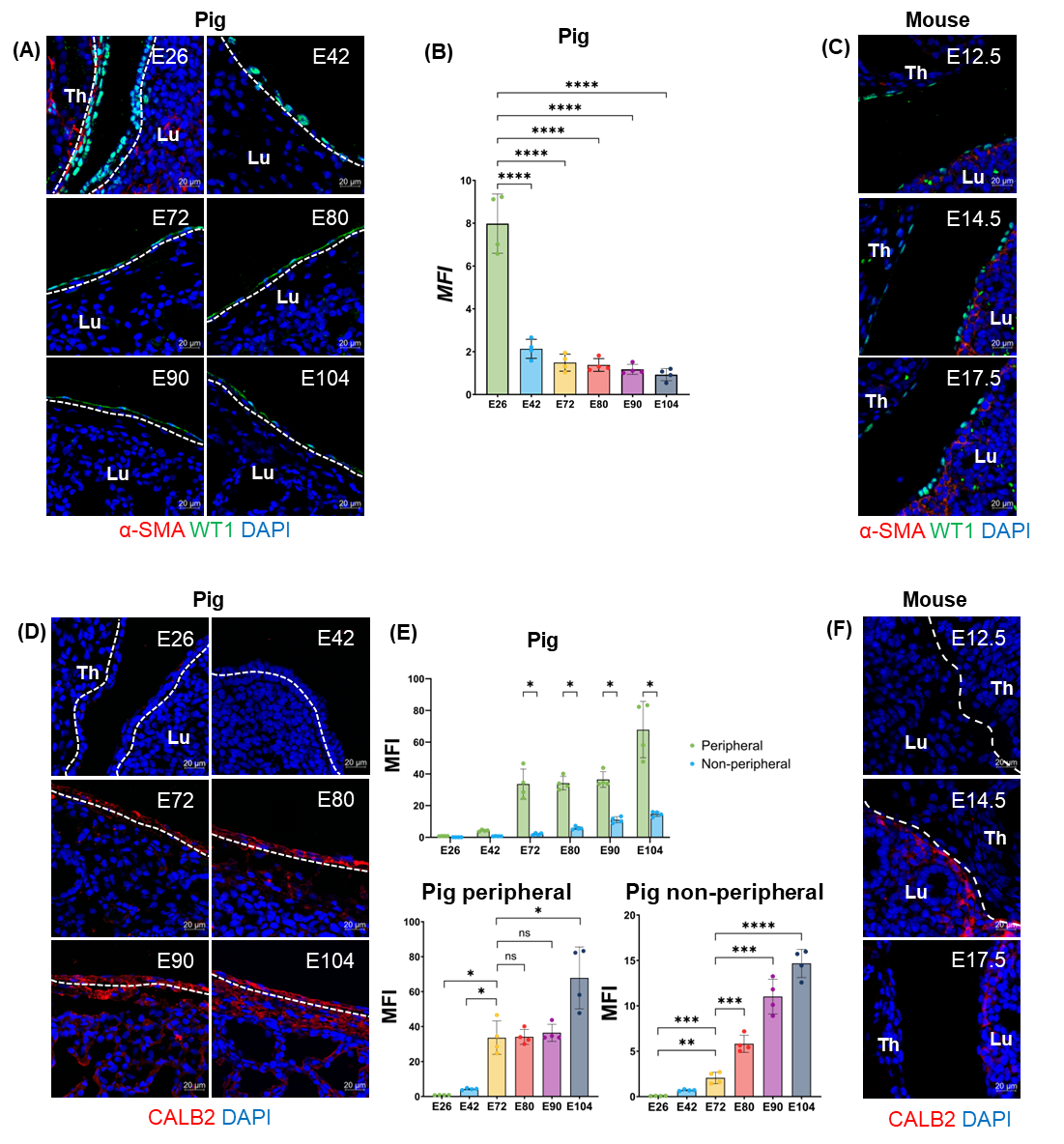

**Figure S5.**

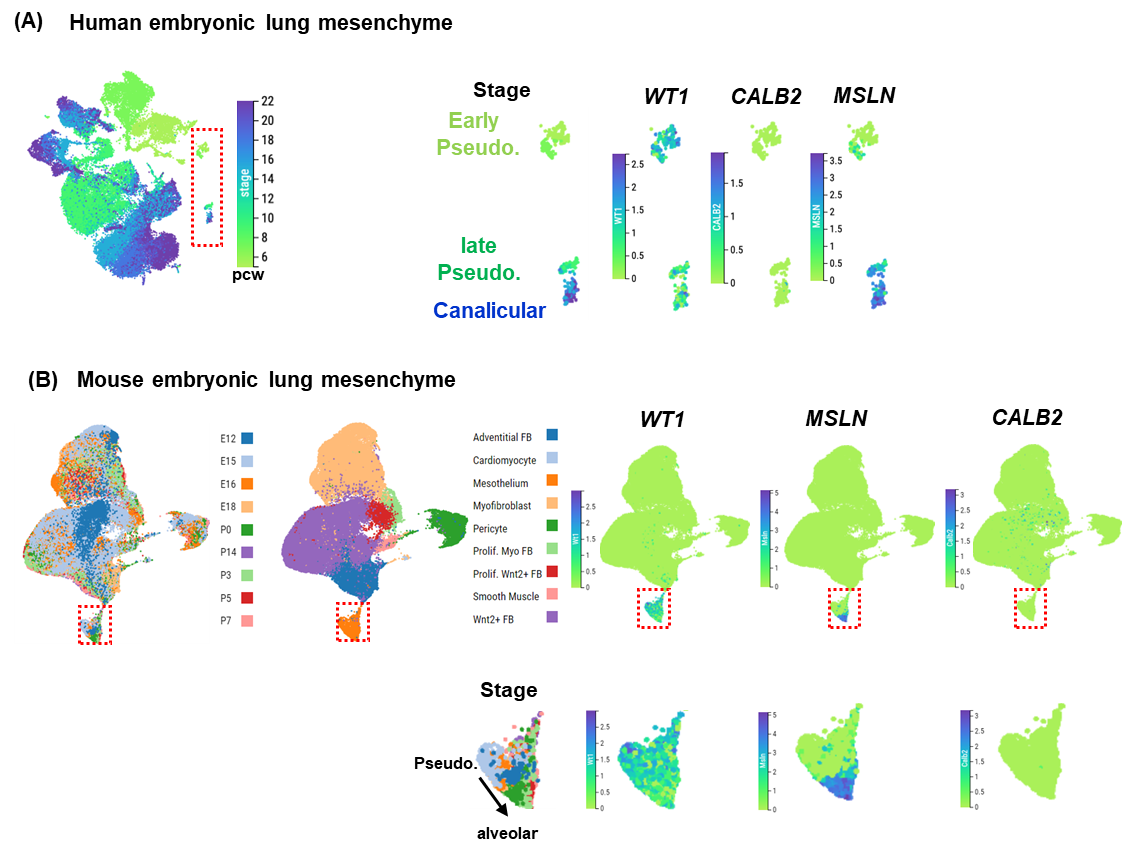

**Figure S6.**

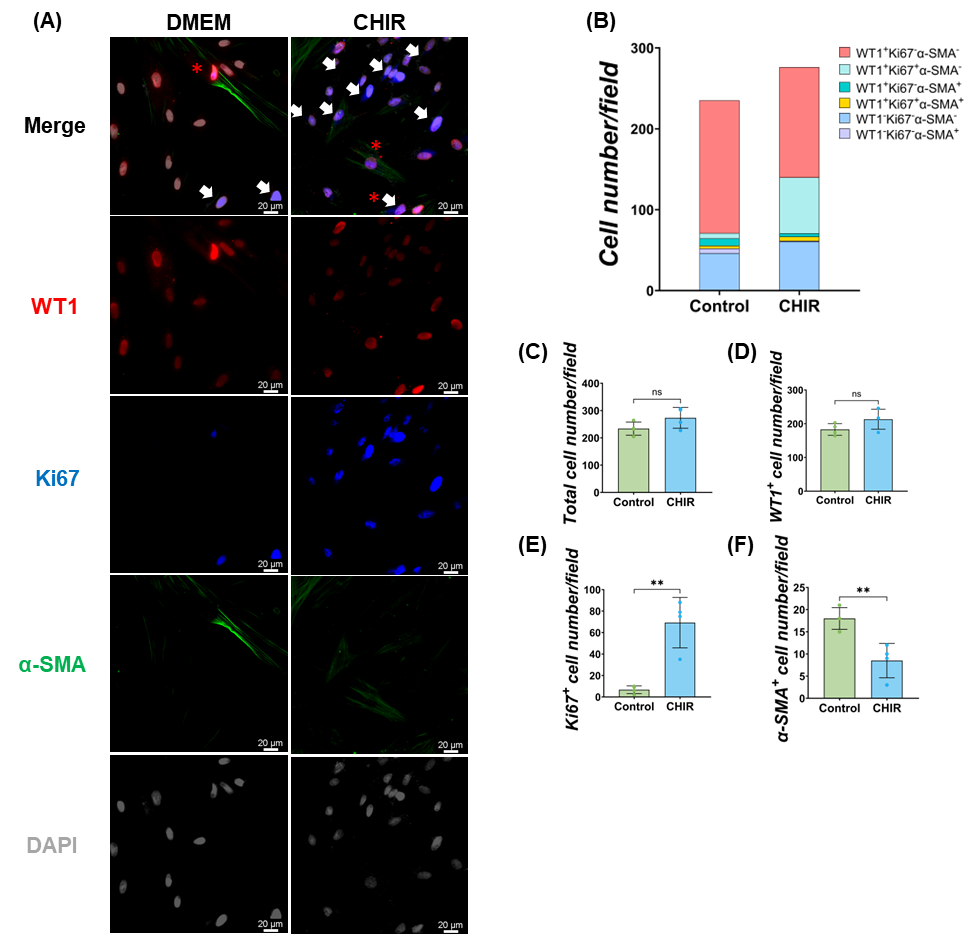

**Table S1.**

| PCR target | Forward primer | Reverse primer |
| --- | --- | --- |
| GAPDH | GTCGGAGTGAACGGATTTGGC | GTGGAGGTCAATGAAGGGGTC |
| WT1 | CTGTCCATTTCTCGGGCCAA | TGACTGTGCTGTATCCCTGGT |
| MSLN | CTCAGCGTTGTTCGGGTGAC | ACCCAGGTAGGGCTTGATCC |
| CALB2 | ATGACAAGAACGCGGATGGG | ACTTCCGCCAAGCCTCCATA |
| α-SMA | ACCGGGAGAAGATGACCCAG | CCAGCACAATGCCAGTCGTA |
| Col1A1 | GTGAAGCTGGTCCCCAAGGA | CAATACCAGGAGCGCCGTTG |
| CD44 | CAGCATCTTCCACACCCACC | AGGGCACAGCTGAAGTAATGGT |
| ITGB1 | CCAAATGGGACACGGGTGAA | TCAGAACCTGCCCATAGCGA |
